# Comparison of aggregation methods for multiphenotype exomic variant prioritization

**DOI:** 10.1101/064899

**Authors:** Alejandro Sifrim, Dusan Popovic, Joris R. Vermeesch, Jan Aerts, Bart De Moor, Yves Moreau

## Abstract

The identification of disease-causing genes in Mendelian disorders has been facilitated by the detection of rare disease-causing variation through exome sequencing experiments. These studies rely on population databases to filter a majority of the putatively neutral variation in the genome and additional filtering steps using either cohorts of diseased individuals or familial information to narrow down the list of candidate variants. Recently, new computational methods have been proposed to prioritize variants by scoring them not only based on their potential impact on protein function but also on their relevance given the available information on the disease under study. Usually these diseases comprise several phenotypic presentations, which are separately prioritized and then aggregated into a global score. In this study we compare several simple (e.g. maximum and mean score) and more complex aggregation methods (e.g. order statistics, parametric modeling) in order to obtain the best possible prioritization performance. We show that all methods perform reasonably well (median rank below 20 out of more than 8000 variants) and that the selection of an optimal aggregation method depends strongly on the fraction of uninformative phenotypes. Finally, we propose guidelines as to how to select an appropriate aggregation method based on knowledge of the phenotype under study.

## Introduction

In recent years, whole-exome sequencing has facilitated the discovery of genes underlying rare Mendelian disease by allowing the identification of disease-causing genomic variation [1–6]. Several different strategies have been proposed to sift through the thousands of variants identified in a single human exome in order to retain a handful of meaningful candidate variants for further downstream functional validation studies[7]. These strategies usually start by filtering variants with a higher potential for disturbing gene function, such as gaining a stop-codon, resulting in a frameshift, changing the amino-acid sequence of the protein or interfering with a splice site. In this first filtering step intergenic, intronic and synonymous mutations are discarded. A second filtering step removes all variants present under a certain minor allele frequency (e.g. the variant occurs in less than 1% of the population) in commonly used population databases such as the 1000 Genomes Project[8] or the NHLBI Exome Sequencing Project[9]. This step relies on the assumption that variants causing rare disease, following either dominant or recessive inheritance patterns, would not be present in high frequencies in populations of healthy individuals. Even after such filtering several hundreds of potential candidate variants remain and additional filtering is required. At this point strategies diverge depending on the available data. If familial samples are available, linkage analysis or searching for *de novo* variants can sufficiently narrow down the list to only a couple of variants[10]. Another strategy is to sequence a number of unrelated individuals exhibiting very similar phenotypical presentations and looking for common genes struck by variants in several of these individuals[2, 3]. This strategy usually assumes a monogenic inheritance pattern in order to have sufficient signal to overcome the noise of neutral genomic variation. Additionally, acquiring sufficient samples is difficult as patient recruitment usually has to happen across several clinical institutions due to the rarity of the disorder in any given population.

In order to assist in this process, several computational methods have been developed to predict the impact of a given variant based on various biochemical, structural and evolutionary properties [11–13]. Although these methods provide reasonable performance in predicting variants affecting protein function, they lack the specificity to identify true disease-causing variants as the human genome is riddled with mildly deleterious variation unrelated to the specific phenotype under study[14–16]. In order to remedy this, new computational approaches were proposed which consider not only functional impact but also phenotypic relevance[17, 18]. In a recent study we proposed a method, named eXtasy, that integrates deleteriousness and haploinsufficiency prediction with phenotype-specific gene prioritization through genomic data fusion[19]. This approach relies on the similarity across several data sources (*e.g.* gene expression, protein-protein interactions, sequence similarity) between known phenotype-associated genes and the mutated gene[20, 21]. This process aids in discriminating variants in genes likely to be involved in the phenotype and variants in genes which might contain functional variants but for which there is no evidence of the genes being involved in the disease.

In this study we elaborate on the problem of aggregating across different prioritizations when the observed clinical presentation is an agglomeration of different phenotypes, as is often the case in rare Mendelian disorders. For example, Miller syndrome, one of the first Mendelian diseases elucidated by massively parallel exome sequencing[3], comprises a combination of craniofacial abnormalities, postaxial limb deformities and sometimes internal malformations. It would be reasonable to assume that some of these phenotypes would be more informative than others when considering them for variant prioritization, as some of them might be better understood in the current literature, they might show less complex known underlying molecular mechanisms or due to the inherent phenotypic variability of the disease. Yet to estimate their relative informativeness would require extremely in-depth knowledge of these properties. To circumvent this problem in eXtasy, we opted to use the maximum score obtained from any single variant prioritization for a given variant. This garantueed that if one of the phenotypes delivered an informative prioritization this would be taken into account (increasing sensitivity of the method), yet potentially also inflated our scores of non-disease causing variants, leading to increased false positive rates. Using the maximum essentially discards information when more than one informative phenotype is used for prioritization as only the best phenotype is considered. In this study we propose 3 alternative ways of aggregating over multiple phenotype-specific prioritizations using either classical order statistics[20, 22], robust rank aggregation[23] and statistical modeling of score distributions. To estimate and compare their performance we also adapt our benchmarking scheme to resemble a more real-life application of variant prioritization. We conclude by showing that although in most cases the maximum outperforms the other aggregation methods, using more complex aggregation methods is usually the best when phenotypes are carefully selected. These methods also offer the added benefit of obtaining probabilistic outcomes which could be integrated further in other statistical frameworks, such as case/control association studies.

## Material and Methods

### Benchmark

In order to compare the performance of the different aggregation methods we set up a benchmark which closely resembles a real-life application of variant prioritization using whole exome sequencing. We first generated 10000 synthetic exomes by randomly sampling nonsynonymous variants from the 1000Genomes Project. For every nonsynonymous variant in the 1000Genomes Project (October 2012 release) a random number between 0 and 2184 (the maximum number of times an allele could be present in the 1092 diploid individuals) was generated and divided by 2184. If this randomly generated number was smaller than the observed global minor allele frequency for that variant, then this variant was included in the synthetic exome. We chose the Human Gene Mutation Database[24] as a set of disease-causing variants, and selected 16272 nonsynonymous variants which were suitable for variant prioritization and studying phenotype aggregation (having at least 3 different phenotypes with at least 3 known gene-phenotype associations each). For each of these variants we randomly selected one of the 10000 synthetic exomes and injected that disease-causing variant into the exome. We then ran these synthetic exomes containing one disease-causing nonsynonymous mutation through eXtasy for each phenotype describing the disease. For each exome we aggregated the resulting scores of the different prioritization either using the maximum, mean, the median, classic and robust order statistics and a statistical modelling approach. We then obtained the rank of the injected disease-causing variant in the synthetic exome for each aggregation method (Figure 1). We additionally computed the fraction of informative phenotypes for a disease-causing variant as the number of phenotypes with an eXtasy score > 0.5 over the the total number of phenotypes prioritized. This threshold is the same at which a variant is considered disease-causing in the original eXtasy paper and corresponds to the majority vote of trees in the Random Forest model.

**Figure 1:**
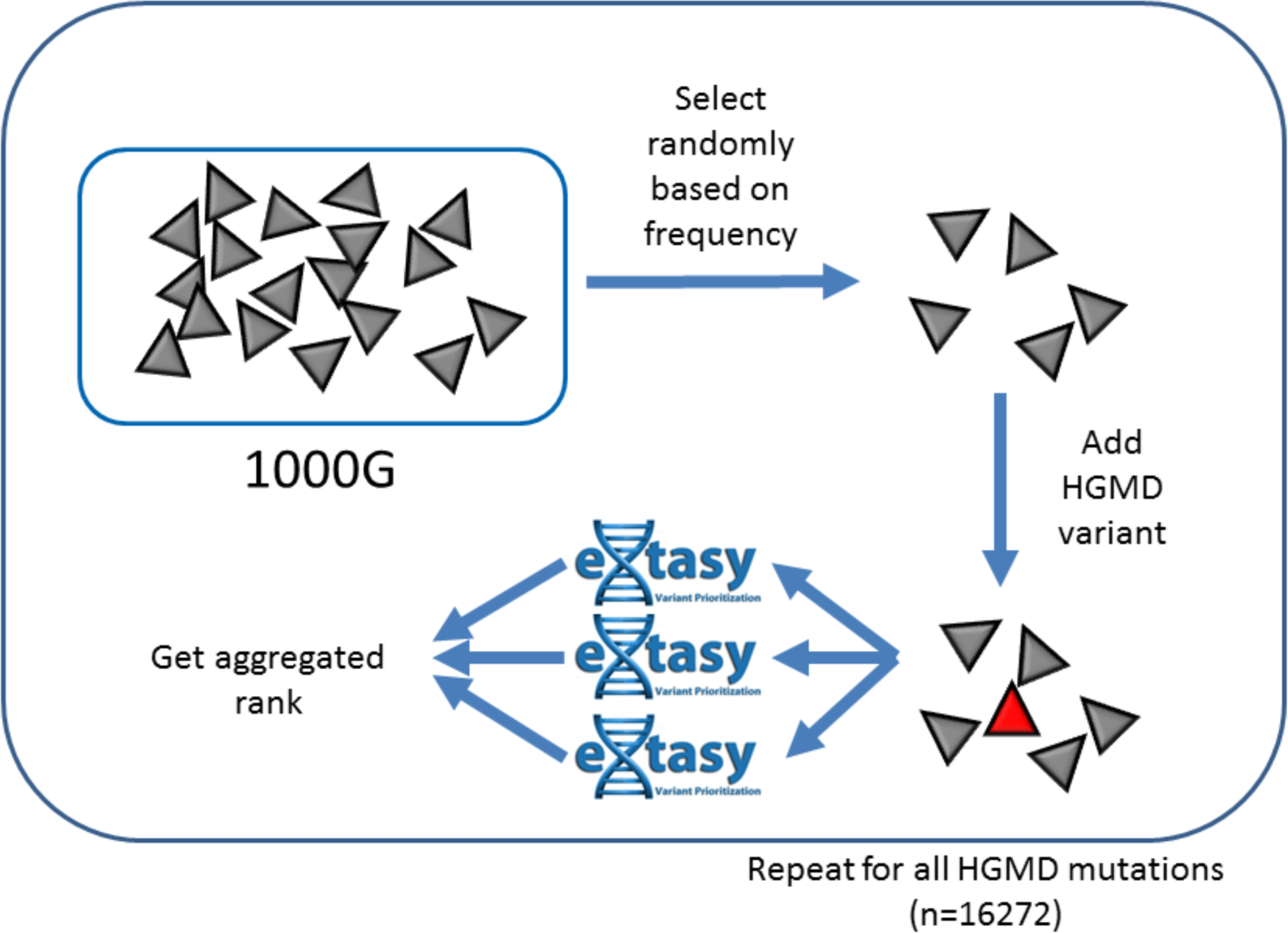
Schematic representation of the benchmarking scheme: Nonsynonymous variants are first sampled from 1000Genomes Project data based on their respective minor allele frequency, generating a synthetic control exome. Then, one disease-causing variant from HGMD is added to a single synthetic control exome and eXtasy prioritizations are performed for each phenotype annotated to the disease-causing variant in that exome. The different phenotype-specific prioritizations are then combined into a single score using different aggregation schemes and the ranks for the injected disease-causing variant is compared across the different methods. This process is repeated for each of the 16272 nonsynonymous HGMD variants.

### Non-parametric Order Statistics

Nonparametric order statistics have been previously used to aggregate rankings of genes obtained from different experiments or data sources[20, 22]. These methods estimate the probability of a given combination of ranks under the null hypothesis that the ranks were randomly drawn. These methods have the benefit if being robust to non-informative rankings, for example adding an erroneous phenotype in the variant prioritization, without losing the added power of additional informative rankings (which are lost by aggregating using the maximum score). In an initial formulation of a gene rank aggregation algorithm, an algorithm was proposed which estimates the cumulative density function of the order statistic under the null hypothesis using a recursive formulation. This algorithm was computationally prohibitive for moderate numbers of ranked lists and was further refined by Aerts et al. (2006) decreasing the algorithmic complexity and thus computational time. Both formulations assume the null hypothesis that all ranked lists are potentially informative. Another computationally efficient formulation called Robust Rank Aggregation (RRA) was proposed[23] where a fraction of the ranked lists is allowed to be uninformative. In their benchmarks they showed that both RRA and the Aerts et al. formulation perform similarly but that the latter algorithm becomes unstable for large numbers of ranked lists (N > 40), but it is unlikely that such numbers of phenotypes would need to be integrated in variant prioritization experiments. In this study we use the R package (RobustRankAggreg) implemented by Kolde et al. which implements both algorithmic order statistic formulations and apply both of them to our benchmark to rank variants according to their estimated significance.

### Parametric Modelling

Order statistic approaches are extremely generalizable as they don’t make any assumptions of the underlying score distributions by transforming the data into rankings. Although this is extremely useful in many situations, such nonparametric approaches are usually less powerful than their parametric counterparts that take these distributions into account. While studying the eXtasy output score distributions we observed highly reproducible non-Gaussian density functions skewed towards 0 and with long tails towards higher scores, which can be best fitted using a Gamma distribution model. In order to come up with an aggregate score across different eXtasy prioritizations we developed a parametrical statistical approach and applied it to our benchmark. Assume *n* prioritizations *N_i_* for *i* = 1..*n* each containing a set of scores *s_i,k_* with *k* = 1..*K* where *K* is the number of variants under study. For each *N_i_* we robustly fitted a Gamma distribution Γ_*i*_ with parameters *k_i_* and *θ_i_* representing the shape and scale of the distribution using the *robust* package in R. We then computed the probability *p_i,k_* of seeing a score equally or more extreme as *s_i,k_* given the cumulative density function of Γ_*i*_. To aggregate these individual probabilities into a single probability of seeing a score as extreme as *s_i,k_* across the *n* different distributions we compute the Fisher’s omnibus meta-analysis statistic 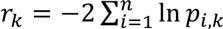 [25]. This allows us to compute the global probability of seeing such a combination of eXtasy scores by computing the probability of *r_k_* given the cumulative density function of a χ^2^ distribution with 2*n* degrees of freedom. Because of the large number of variants tested we correct these probabilities using Benjamini-Hochberg False Discovery Rate correction. We then rank the variants for each synthetic exome in our benchmark given the aggregated multiple testing corrected p-value.

## Results

In contrast with the previously reported eXtasy case-control classification benchmark, our goal with the synthetic exome benchmark was to simulate a real-life situations where variant prioritization would be used and resembles the validation benchmark proposed by Robinson et al. [18]. Our exome generation algorithm generated exomes containing an average of 8399 nonsynonymous variants (n=10000, sd=242). This number is higher than the roughly 6000 variants reported from large-scale exome sequencing projects[9], but is in line with whole-genome sequencing studies[8, 26] which do not face the same difficulties that exome-capturing techniques due to coverage variability. As an indication of real-life performance we describe the rank of the injected disease-causing variant as a performance metric, as opposed to classical metrics (e.g. precision/recall), to compare aggregation schemes for multiphenotype variant prioritization using eXtasy.

Overall, we observe that all aggregation methods perform reasonably well (Table 1), having a median rank for the disease-causing mutation in the top 20 and a below the top 100 in 75% of the cases (with the exception of the median eXtasy score and RRA). Looking at the cumulative distribution functions of the ranks of the different methods (Figure 2), we can see that using the maximum eXtasy score (as used in the original eXtasy publication) delivers the best rankings compared to the other methods. Aggregation using the average eXtasy score or the more involved parametric and nonparametric methods seem to show similar performance. Finally, RRA and the median eXtasy score provide the worst performance out of all the aggregation schemes. These results indicate that uninformative or incorrectly prioritized phenotypes might affect the global parameters ranking significantly, which might explain why using the maximum score (being the most robust method) generally outperforms other aggregation schemes. In order to further investigate this hypothesis, we looked at how varying degrees of uninformative phenotypes affect the different aggregation schemes by stratifying our benchmark according to the fraction of individual phenotypes correctly prioritized.

**Figure 2:**
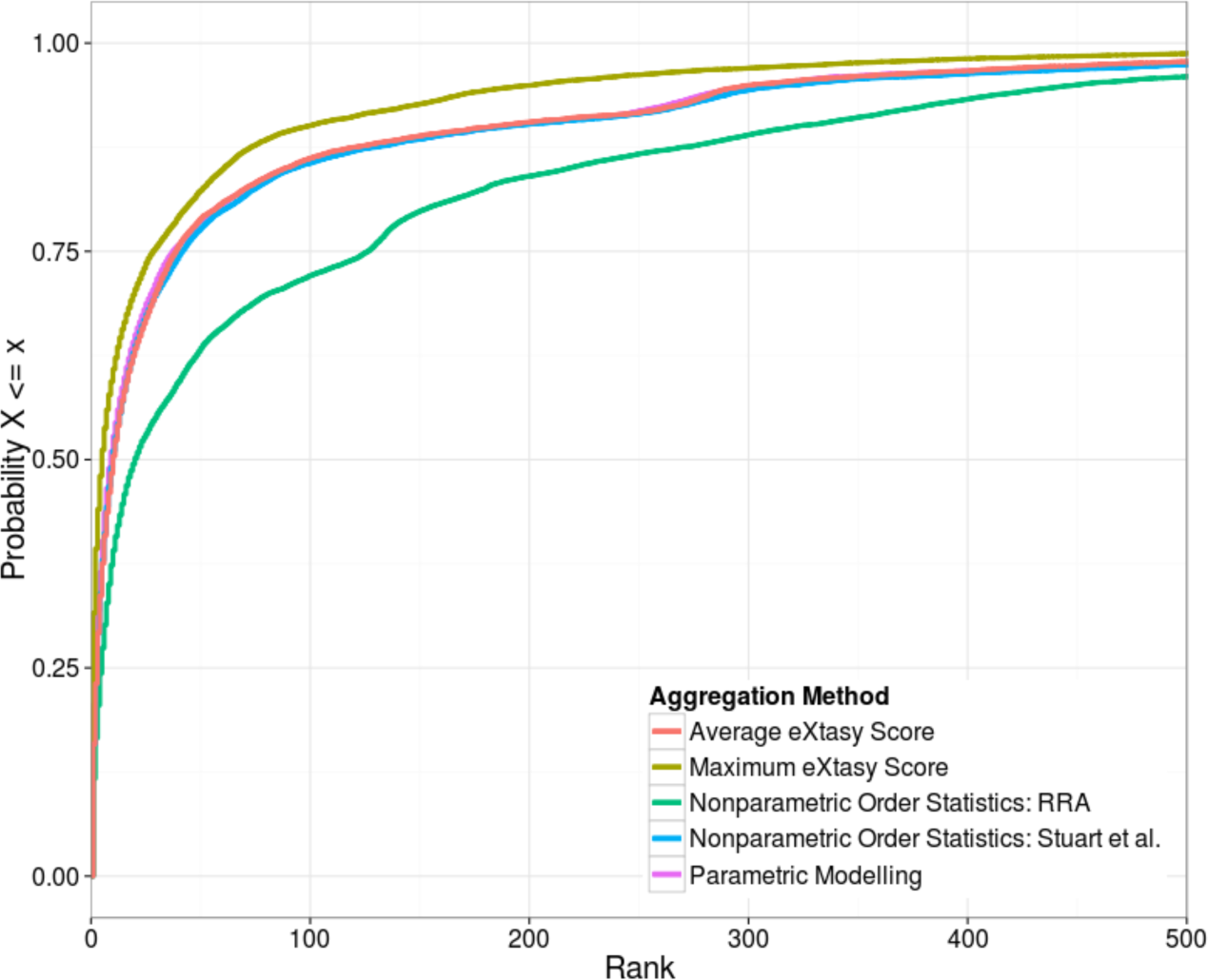
Empirical Cumulative Distribution Function (ECDF) of ranks comparing different aggregation schemes: This chart shows the proportion of variants out of all disease-causing variants (n=16272) with global ranks after aggregation lower or equal than the rank shown on the x-axis.

**Table 1:** Summary statistics for the overall comparison of different aggregation methods: This table shows the mean, median, 25%-quantile (Q1) and 75%-quantile of the global ranks of disease-causing variants (n=16272) using different aggregation schemes.

| Aggregation Method | Q1 | Median | Mean | Q3 |
| --- | --- | --- | --- | --- |
| Average eXtasy Score | 3 | 10 | 59 | 39 |
| Maximum eXtasy Score | 1 | 5 | 41 | 28 |
| Nonparametric Order Statistics: Stuart et al. | 3 | 10 | 64 | 42 |
| Nonparametric Order Statistics: RRA | 5 | 20 | 102 | 126 |
| Parametric Modelling | 2 | 9 | 111 | 37 |

After stratification into low (n=5862), moderate (n=8559) and high (n=1837) fractions of informative phenotypes we observe that no single aggregation method performs best across the three classes (Figure 3). When only less than a third of phenotypes are informative, the average rank of the disease-causing variants degrades significantly independently of the aggregation method (Table 2). In this scenario, aggregation using the maximum eXtasy score delivers the best ranking compared to all other methods, while all other methods show nearly equal performance (with the exception of RRA, showing significantly lower performance). When roughly halve the phenotypes are informative, all aggregation methods (except RRA) perform reasonably well (average rank lower in 10, and 75% of variants ranked below 12). Additionally, methods which leverage information across different phenotypes (mean score, order statistics, parametric modelling) perform slightly better than aggregation using the maximum score. In the case where more than two-thirds of the phenotypes are informative 75% of the disease-causing variants are in the top 3 ranked mutations for all methods except aggregation by maximum score (75% of variants in the top 10).

**Figure 3:**
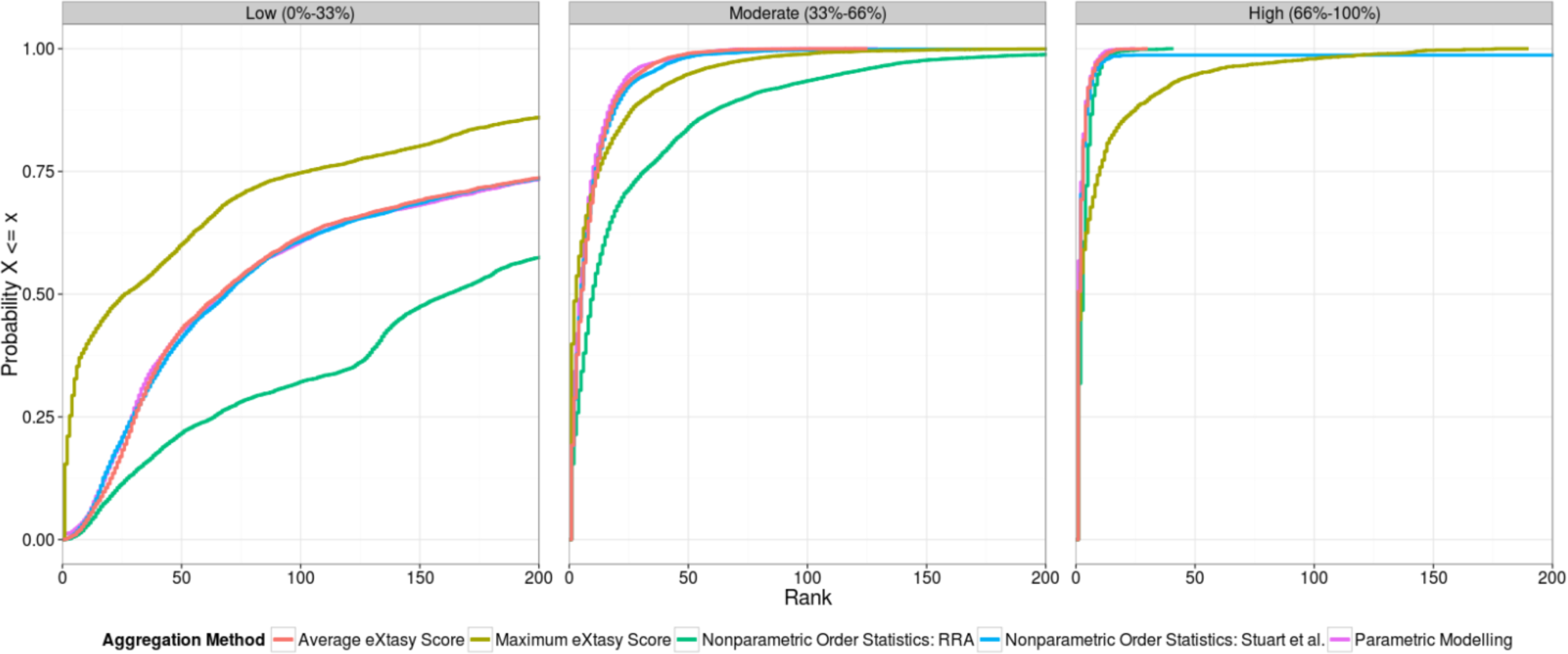
Empirical Cumulative Distribution Function (ECDF) of ranks by fraction of informative phenotypes comparing different aggregation schemes: This chart shows the proportion of variants out of all disease-causing variants for fractions of informative phenotypes divided into 3 classes: Low (n=5862), Moderate (n=8559) and High (n=1837) with respectively 0%-33%, 33%-66% and 66%-100% of phenotypes showing an eXtasy score > 0.5 for the prioritized disease-causing variant.

**Table 2:** Summary statistics for the comparison of different aggregation schemes by fraction of informative phenotypes: Analogous to Table 1, this table shows summary statistics for the ranks of disease-causing mutation but after stratification into 3 classes representing different fractions of informative phenotypes: low (0%-33%), moderate (33%-66%) and high (66%-100%) number of informative phenotypes showing an eXtasy score > 0.5 for the prioritized disease-causing variant.

| Class | Aggregation Method | Q1 | Median | Mean | Q3 |
| --- | --- | --- | --- | --- | --- |
| Low<br>(0%-33%) | Average eXtasy Score | 31 | 66 | 152 | 218 |
|  | Maximum eXtasy Score | 3 | 26 | 95 | 102 |
|  | Nonparametric Order Statistics: Stuart et al. | 30 | 69 | 158 | 224 |
|  | Nonparametric Order Statistics: RRA | 64 | 162 | 243 | 347 |
|  | Parametric Modelling | 29 | 67 | 295 | 222 |
| Moderate<br>(33%-66%) | Average eXtasy Score | 2 | 5 | 8 | 12 |
|  | Maximum eXtasy Score | 1 | 3 | 11 | 12 |
|  | Nonparametric Order Statistics: Stuart et al. | 2 | 5 | 10 | 11 |
|  | Nonparametric Order Statistics: RRA | 3 | 10 | 26 | 31 |
|  | Parametric Modelling | 1 | 5 | 8 | 10 |
| High<br>(66%-100%) | Average eXtasy Score | 1 | 1 | 2 | 3 |
|  | Maximum eXtasy Score | 1 | 2 | 11 | 10 |
|  | Nonparametric Order Statistics: Stuart et al. | 1 | 1 | 11 | 3 |
|  | Nonparametric Order Statistics: RRA | 1 | 3 | 3 | 5 |
|  | Parametric Modelling | 1 | 1 | 2 | 3 |

## Discussion

In this study we investigate the performance of different aggregation schemes when dealing with multiphenotype exomic variant prioritization. We benchmark these methods mimicking a real-life application by generating synthetic healthy exomes and injecting single known disease-causing nonsynonymous mutations from HGMD. By comparing different aggregation schemes, ranging from simple metrics to more involved nonparametric and parametric methods, we show that no single method is optimal for all cases and that performance depends on the expected fraction of informative phenotypes.

Not unsurprisingly, the optimal aggregation method is dependent on the signal-to-noise ratio originating from the underlying individual prioritizations. Any prioritization for a specific phenotype can add noise to the global prioritization for two reasons. First, the specific phenotype might not be biologically well-suited to describe the disease under study. This can for example be attributed to phenotypic variability of the disease (e.g. some of the phenotypes might not always be present in all patients showing the disease). This problem can relatively easily be managed by clinical knowledge of the disease, and thus be avoided in real-life applications. In our benchmark this is more difficult to address due to the sheer number of included diseases/phenotypes where disease terms are linked to phenotypic ontologies automatically. Secondly, the gene prioritization step can be noisy due to the selected set of training genes. For example, certain phenotypes could be overly general (e.g. intellectual disability, abnormality of the face) and have many heterogeneous underlying molecular mechanisms. Using all known genes in the gene prioritization step for such phenotypes results in uninformative prioritizations due to a dilution of the signal where similarity is estimated to an extremely diverse group of genes and thus leads to reduced performance. The inverse problem also exists where too little is known about the phenotype, biasing the prioritization towards certain training genes which might not necessarily be similar to the as-of-yet undiscovered causative gene. Predicting *a priori* the performance of the gene prioritization is difficult and requires a good understanding of the underlying data being used.

Because of these problems, it is important to use aggregation methods which can handle these uninformative or erroneous prioritizations robustly. Aggregating using the maximum score is a simple and intuitive robust solution and performs well in our previous and current benchmark, especially in low signal-to-noise situations. The drawback of using the maximum is that it ignores information from additional phenotypes, which could be quite powerful in increasing the performance of a global prioritization. This is indicated by the better performance of methods which take advantage of these phenotypes (mean, order statistics and parametric modelling) when better signal-to-noise ratios are observed. It is important to remark that uninformative phenotypes are likely overrepresented in our benchmark, compared to real-life applications, because of the automatic mappings generated between disease-variants and phenotypic ontologies and the lack of clinical expertise for each evaluated disease. Considering this, it is likely that using these methods will result in better variant prioritizations if the phenotypes are carefully selected.

Here we show that choosing a suitable aggregation method in multiphenotype variant prioritization can affect the results of the resulting global prioritization. We demonstrate this by using simple aggregation methods such as the maximum score and the mean score. Additionally we show that more involved nonparametric and parametric statistical methods offer similar performance with the added benefit of providing estimates of significance which could in future studies could easily be used in broader statistical frameworks such as those used in weighted association or familial studies [27, 28]. We do remark that these methods assume independence of the individual prioritizations, an assumption which is likely not true due to the usage of phenotype-aspecific data sources in eXtasy. This might not greatly influence their performance in a ranking scenario but result in biased estimates of significance. Several methods have been proposed which handle these dependencies [29, 30]but were not considered in this study due to the unavailability of usable implementations, but should be considered when true statistical significance is of importance.

Although currently we rely on the user to select an appropriate aggregation method given his experience on how well-characterised the phenotype under study is. In future work we hope to partially or fully automate this process by trying to evaluate training gene set heterogeneity and gene prioritization performance through cross-validation. By gradually improving variant prioritization performance and usability we hope to aid clinical geneticists in discovering disease-causing mutations for their respective phenotypes of interest.

We currently offer all the proposed aggregation schemes in the current version of eXtasy (http://homes.esat.kuleuven.be/∼bioiuser/eXtasy/). We also publicly provide the benchmark data for further development or benchmarking of other score/rank aggregation methodology (http://homes.esat.kuleuven.be/∼bioiuser/eXtasy/aggregation.tar.gz).

## Reference

1. Hoischen A, van Bon BWM, Gilissen C, Arts P, van Lier B, et al. (2010) De novo mutations of SETBP1 cause Schinzel-Giedion syndrome. Nat Genet 42: 483–485. Available: http://www.ncbi.nlm.nih.gov/pubmed/20436468. Accessed 26 April 2011.

2. Van Houdt JKJ, Nowakowska BA, Sousa SB, van Schaik BDC, Seuntjens E, et al. (2012) Heterozygous missense mutations in SMARCA2 cause Nicolaides-Baraitser syndrome. Nat Genet 44: 445–449. Available: http://www.ncbi.nlm.nih.gov/pubmed/22366787. Accessed 29 March 2012.

3. Ng SB, Buckingham KJ, Lee C, Bigham AW, Tabor HK, et al. (2010) Exome sequencing identifies the cause of a mendelian disorder. Nat Genet 42: 30–35. Available: http://www.pubmedcentral.nih.gov/articlerender.fcgi?artid=2847889&tool=pmcentrez&rendertype=abstract. Accessed 17 August 2010.

4. Lyon G, Wang K(2012) Identifying disease mutations in genomic medicine settings: current challenges and how to accelerate progress. Genome Med. Available: http://www.biomedcentral.com/content/pdf/gm359.pdf. Accessed 30 August 2012.

5. Choi M, Scholl UI, Ji W, Liu T, Tikhonova IR, et al. (2009) Genetic diagnosis by whole exome capture and massively parallel DNA sequencing. Proc Natl Acad Sci U S A 106: 19096–19101. Available: /pmc/articles/PMC2768590/?report=abstract. Accessed 15 December 2013.

6. Ng SB, Bigham AW, Buckingham KJ, Hannibal MC, McMillin MJ, et al. (2010) Exome sequencing identifies MLL2 mutations as a cause of Kabuki syndrome. Nat Genet 42: 790–793. Available: /pmc/articles/PMC2930028/?report=abstract. Accessed 16 December 2013.

7. Gilissen C, Hoischen A, Brunner HG, Veltman JA (2012) Disease gene identification strategies for exome sequencing. Eur J Hum Genet 20: 490–497. Available: /pmc/articles/PMC3330229/?report=abstract. Accessed 12 December 2013.

8. Durbin RM, Altshuler DL, Abecasis GR, Bentley DR, Chakravarti A, et al. (2010) A map of human genome variation from population-scale sequencing. Nature 467: 1061–1073. Available: http://www.nature.com/doifinder/10.1038/nature09534.Accessed 27 October 2010.

9. Tennessen JA, Bigham AW, O’Connor TD, Fu W, Kenny EE, et al. (2012) Evolution and functional impact of rare coding variation from deep sequencing of human exomes. Science (80-) 337: 64–69. Available: http://www.ncbi.nlm.nih.gov/pubmed/22604720. Accessed 12 February 2013.

10. Vissers LELM, de Ligt J, Gilissen C, Janssen I, Steehouwer M, et al. (2010) A de novo paradigm for mental retardation. Nat Genet 42: 1109–1112. Available: http://www.ncbi.nlm.nih.gov/pubmed/21076407. Accessed 30 May 2011.

11. Ng P, Henikoff S (2003) SIFT: Predicting amino acid changes that affect protein function. Nucleic Acids Res 31: 3812. Available:http://nar.oxfordjournals.org/content/31/13/3812.short. Accessed 30 May 2011.

12. Schwarz JM, Rödelsperger C, Schuelke M, Seelow D (2010) MutationTaster evaluates disease-causing potential of sequence alterations. Nat Methods 7: 575–576. Available: http://www.ncbi.nlm.nih.gov/pubmed/20676075. Accessed 13 July 2011.

13. Adzhubei IA, Schmidt S, Peshkin L, Ramensky VE, Gerasimova A, et al. (2010) A method and server for predicting damaging missense mutations. Nat Methods 7: 248–249. Available:http://www.pubmedcentral.nih.gov/articlerender.fcgi?artid=2855889&tool=pmcentrez&rendertype=abstract. Accessed 15 April 2011.

14. Kumar S, Sanderford M, Gray VE, Ye J, Liu L (2012) Evolutionary diagnosis method for variants in personal exomes. Nat Methods 9: 855–856. Available: http://www.ncbi.nlm.nih.gov/pubmed/22936163. Accessed 19 February 2013.

15. Kryukov G V, Pennacchio LA, Sunyaev SR (2007) Most rare missense alleles are deleterious in humans: implications for complex disease and association studies. Am J Hum Genet 80: 727–739. Available: http://www.pubmedcentral.nih.gov/articlerender.fcgi?artid=1852724&tool=pmcentrez&rendertype=abstract. Accessed 13 December 2013.

16. O’Fallon B, Wooderchak-Donahue W, Bayrak-Toydemir P, Crockett D (2013) VarRanker: rapid prioritization of sequence variations associated with human disease. BMC Bioinformatics 14: S1. Available: http://www.biomedcentral.com/1471-2105/14/S13/S1. Accessed 18 October 2013.

17. Moreau Y, Tranchevent L-C (2012) Computational tools for prioritizing candidate genes: boosting disease gene discovery. Nat Rev Genet 13: 523–536. Available: http://www.ncbi.nlm.nih.gov/pubmed/22751426. Accessed 12 July 2012.

18. Robinson P, Köhler S, Oellrich A, Wang K, Mungall C, et al. (2013) Improved exome prioritization of disease genes through cross species phenotype comparison. Genome Res: gr.160325.113–. Available: http://genome.cshlp.org/content/early/2013/10/25/gr.160325.113.abstract?cited-by=yes&legid=genome;gr.160325.113v1. Accessed 12 December 2013.

19. Sifrim A, Popovic D, Tranchevent L-C, Ardeshirdavani A, Sakai R, et al. (2013) eXtasy: variant prioritization by genomic data fusion. Nat Methods. Available: http://dx.doi.org/10.1038/nmeth.2656. Accessed 1 October 2013.

20. Aerts S, Lambrechts D, Maity S, Van Loo P, Coessens B, et al. (2006) Gene prioritization through genomic data fusion. Nat Biotechnol 24: 537–544. Available: http://www.ncbi.nlm.nih.gov/pubmed/16680138.

21. Tranchevent L-C, Barriot R, Yu S, Van Vooren S, Van Loo P, et al. (2008) ENDEAVOUR update: a web resource for gene prioritization in multiple species. Nucleic Acids Res 36: W377–84. Available: http://www.ncbi.nlm.nih.gov/pubmed/18508807.

22. Stuart J, Segal E, Koller D, Kim S (2003) A gene-coexpression network for global discovery of conserved genetic modules. Science (80-). Available: http://www.sciencemag.org/cgi/content/full/302/5643/249.

23. Kolde R, Laur S, Adler P, Vilo J (2012) Robust rank aggregation for gene list integration and meta-analysis. Bioinformatics 28: 573–580. Available: http://bioinformatics.oxfordjournals.org/content/28/4/573.short. Accessed 22 December 2013.

24. Stenson PD, Mort M, Ball E V, Howells K, Phillips AD, et al. (2009) The Human Gene Mutation Database: 2008 update. Genome Med 1: 13. Available: http://www.biomedcentral.com/content/pdf/gm13.pdf. Accessed 19 February 2013.

25. Fisher R (1925) Statistical methods for research workers. Available: http://scholar.google.be/scholar?hl=en&q=Fisher%2C+R.A.+%281925%29.+Statistical+Methods+for+Research+Workers&btnG=&as_sdt=1%2C5&as_sdtp=#1. Accessed 22 December 2013.

26. Lupski JR, Reid JG, Gonzaga-Jauregui C, Rio Deiros D, Chen DCY, et al. (2010) Whole-genome sequencing in a patient with Charcot-Marie-Tooth neuropathy. N Engl J Med 362: 1181–1191. Available: http://www.ncbi.nlm.nih.gov/pubmed/20220177. Accessed 7 October 2011.

27. Rope AF, Wang K, Evjenth R, Xing J, Johnston JJ, et al. (2011) Using VAAST to identify an X-linked disorder resulting in lethality in male infants due to N-terminal acetyltransferase deficiency. Am J Hum Genet 89: 28–43. Available: http://www.pubmedcentral.nih.gov/articlerender.fcgi?artid=3135802&tool=pmcentrez&rendertype=abstract. Accessed 5 March 2012.

28. Ionita-Laza I, Makarov V, Yoon S, Raby B, Buxbaum J, et al. (2011) Finding disease variants in Mendelian disorders by using sequence data: methods and applications. Am J Hum Genet 89: 701–712. Available: http://www.pubmedcentral.nih.gov/articlerender.fcgi?artid=3234377&tool=pmcentrez&rendertype=abstract. Accessed 17 March 2012.

29. Kost JT, McDermott MP (2002) Combining dependent P-values. Stat Probab Lett 60: 183–190. Available: http://www.sciencedirect.com/science/article/pii/S0167715202003103. Accessed 22 December 2013.

30. Brown M (1975) 400: A Method for Combining Non-Independent, One-Sided Tests of Significance. Biometrics. Available: http://www.jstor.org/stable/10.2307/2529826. Accessed 22 December 2013.

